# The intrinsically disordered tail of ESCO1 binds DNA in a charge-dependent manner

**DOI:** 10.1101/2023.12.05.570177

**Authors:** Jeffrey R. Schoen, Jingrong Chen, Susannah Rankin

## Abstract

ESCO1 is an acetyltransferase enzyme that regulates chromosome organization and gene expression. It does this by modifying the Smc3 subunit of the Cohesin complex. Although ESCO1 is enriched at the base of chromatin loops in a Cohesin-dependent manner, precisely how it interacts with chromatin is unknown. Here we show that the basic and intrinsically disordered tail of ESCO1 binds DNA with very high affinity, likely through electrostatic interaction. We show that neutralization of positive residues in the N-tail reduces both DNA binding in vitro and association of the enzyme with chromatin in cells. Additionally, disruption of the chromatin state and charge distribution reduces chromatin bound ESCO1. Strikingly, defects in DNA binding do not affect total SMC3 acetylation or sister chromatid cohesion, suggesting that ESCO1-dependent acetylation can occur independently of direct chromatin association. We conclude that the intrinsically disordered tail of ESCO1 binds DNA with both high affinity and turnover, but surprisingly, ESCO1 catalytic activity occurs independently of direct DNA binding by the enzyme.

## Introduction

The Cohesin complex tethers sister chromatids together and organizes chromatin into loops and domains (1–3). These distinct activities of Cohesin are regulated by acetylation of the SMC3 subunit of cohesin, which promotes retention of the complex on chromatin (4–9). In vertebrates, the majority of Cohesin acetylation is mediated by ESCO1, an enzyme which both regulates chromatin organization and contributes to sister chromatid cohesion through Cohesin acetylation (8, 10–12).

Both ESCO1 and its homolog ESCO2 bind chromosomes through their distinct N-terminal tails (13). These N-terminal tails direct their distinct functions (10). ESCO2 interacts with multiple DNA replication proteins through its tail which drives it onto chromatin during S phase (14–17). ESCO1, unlike its homolog, remains associated with chromatin throughout the cell cycle (11, 13). The identity of the direct interacting partner of the ESCO1 N-terminal tail on chromatin remains a mystery.

To better understand ESCO1 function, we have investigated how the protein interacts with chromatin through its N-terminal tail. This region of the protein is rich in basic amino acid residues, and thus has an elevated isoelectric point (pI). By building a series of increasingly neutralized N-terminal tail mutants, we show that interaction of the ESCO1 tail with chromatin is dependent upon its charge. Moreover, we find that ESCO1 is able to bind directly to DNA in vitro through its N-terminal tail in a DNA sequence-independent manner. Surprisingly, ESCO1-dependent Cohesin acetylation and sister chromatid cohesion were not disrupted by loss of this DNA binding tail, indicating that acetylation of Cohesin occurs independent of ESCO1’s direct DNA binding.

## Results

### The N-terminal tail of ESCO1 binds chromatin in a DNA-dependent manner

Although genomic localization of ESCO1 in human cells overlaps with Cohesin, the bulk association of ESCO1 with chromatin is unaffected by loss of Cohesin (9, 18). The N-terminal tail of ESCO1 was previously shown to promote its interaction with chromatin (13). This region contains no known Cohesin-interaction sites; the catalytic domain and a Pds5 (a cohesin-associated protein) binding site are located elsewhere in the protein (18, 19).

To confirm the previously-reported dependence of chromatin association of ESCO1 on its N-terminal tail, we isolated the insoluble nuclear fraction from HeLa cells lacking endogenous ESCO1 and expressing flag epitope-tagged ESCO1 WT or an N-terminal tail deletion mutant (Δ1-200) (10). Immunoblot analysis of the isolated chromatin recapitulated the earlier finding that deletion of the N-terminal 200 amino acids of ESCO1 makes it significantly more soluble (Fig. 1A and B).

**Figure 1.**
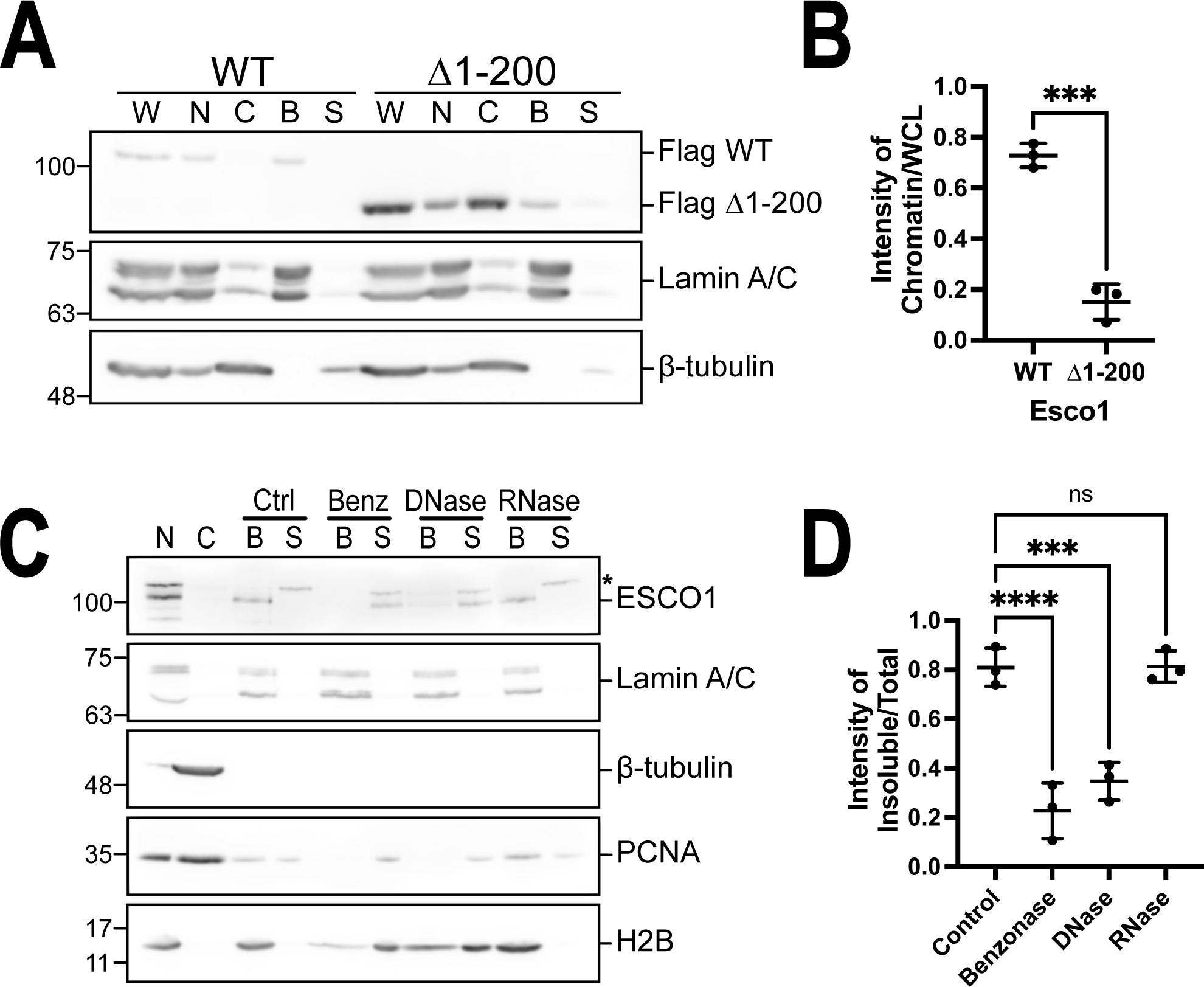
ESCO1 chromatin binding. **A. N-terminus-dependent chromatin association.** HeLa cells expressing Flag-tagged wild-type or N-terminal deleted (Δ1-200) ESCO1 were fractionated to assess ESCO1 chromatin binding by immunoblot analysis. W: whole cell lysate, C: Cytoplasm, B: Bound nuclear fraction, S: Supernatant. **B**. **Quantification:** The experiment shown in A was repeated 3 times and the fraction of Flag-ESCO1 remaining chromatin bound was assessed (*** = p < 0.001; t-test). **C**. **DNA-dependent chromatin association of ESCO1.** Nuclei isolated from 293T cells were subjected to enzymatic digestion as indicated, nuclei were reisolated, and bound and released proteins were assessed by immunoblot as shown. N: Nuclei, C: Cytoplasm, B: Bound nuclear fraction, S: Supernatant. **D**. **Quantification:** The experiment shown in (C) was repeated 3 times and the fraction of ESCO1 remaining nuclear associated was determined (**** = p < 0.0001; *** = p < 0.001; one way ANOVA with DunneX’s correction for multiple comparisons).

To further investigate the nature of ESCO1 binding to chromatin, we treated isolated nuclei to determine whether ESCO1 chromatin binding is dependent on DNA or RNA. Using the non-specific nuclease Benzonase (for both DNA and RNA degradation), DNase I (for DNA degradation), or RNase A (for RNA degradation) we found that ESCO1 retention in nuclei is critically dependent upon the presence of nuclear DNA. Digestion with Benzonase and DNase, but not RNase, released ESCO1 from isolated nuclei (Fig. 1C and 1D). The chromatin-associated proteins PCNA, the core histone H2B, and nuclear envelope associated lamins were used as controls to confirm effectiveness of the extraction protocols. We conclude that chromatin association of endogenous ESCO1 depends on the presence of nuclear DNA.

### Positively charged residues in the ESCO1 N-terminal tail promote chromatin association

We next sought to clarify how the N-terminal tail of ESCO1 promotes its association with chromatin throughout the cell cycle. Crystal structures of ESCO1 show a conserved structured catalytic domain at the C-terminus but have so far have failed to capture its N-terminal tail (20, 21). The AlphaFold prediction model fails to predict structure of this region of the protein and indicates a high degree of disorder throughout the N-terminal tail (22).

The N terminal ∼200 amino acid region of ESCO1 is poorly conserved at the primary sequence level (Table 1). We did, however, note a high distribution of positive residues in this region, resulting in a high average pI (mean pI 10.8). We also noted, consistent with the AlphaFold analysis, that this region of the protein is predicated to be highly disordered (Fig.2A). Charge has been shown to promote the interaction of proteins with condensed mitotic chromosomes (23). This is particularly interesting because while ESCO1 colocalizes with Cohesin throughout interphase, it remains bound to mitotic chromosomes when Cohesin is largely removed (9, 10, 13, 18, 24). Therefore, we hypothesized that ESCO1 could be directly interacting with chromatin through its positively charged and disordered N-terminal tail.

**Figure 2.**
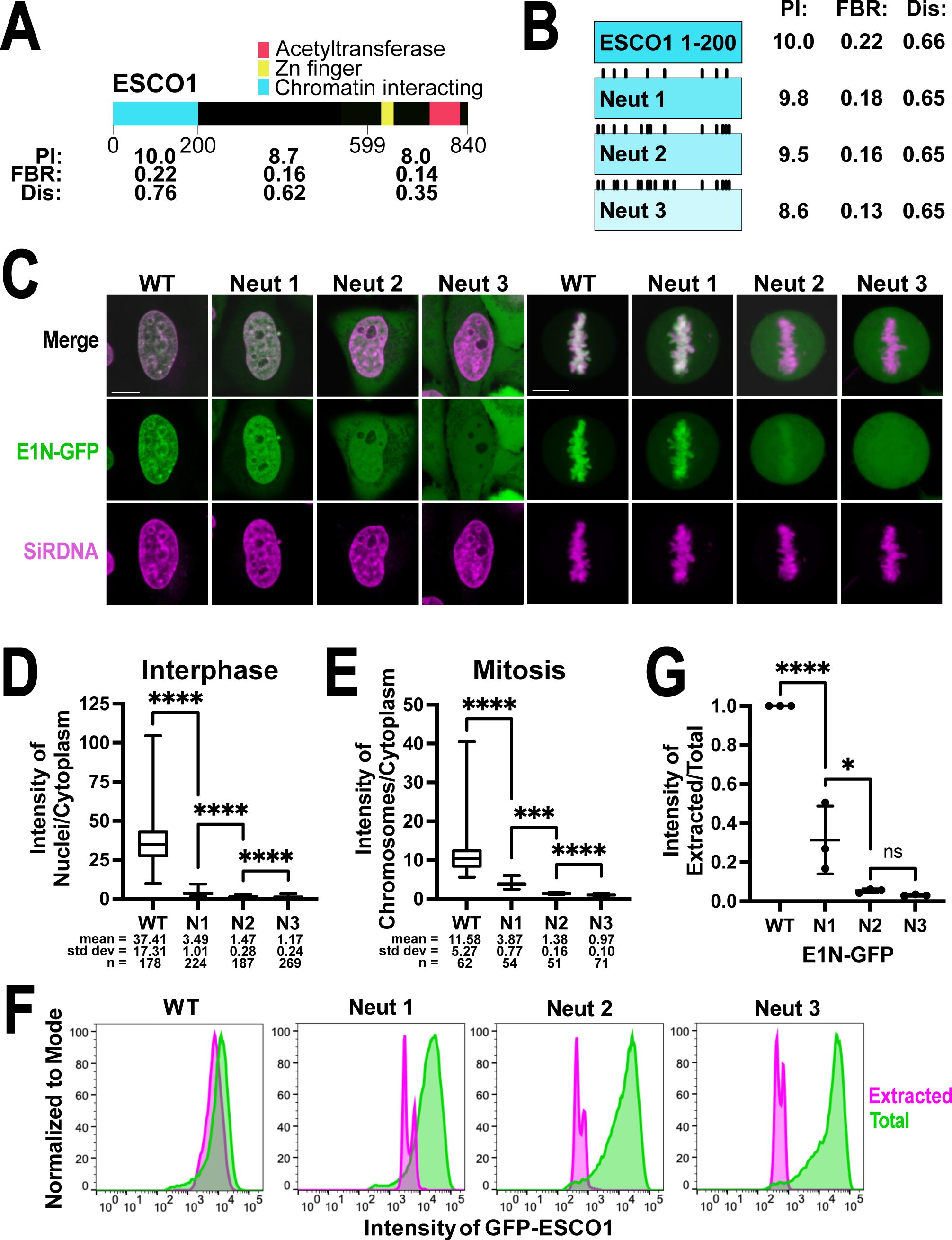
Positive charge promotes association of ESCO1 with chromatin. A. The ESCO1 N-terminus is poorly conserved, disordered, and basic. Diagram of human ESCO1 showing the PI, fraction of basic residues (FBR), and propensity for disorder calculated by PrDOS (0-0.5 = ordered, 0.5-1 = disordered) (54). The chromatin interacting domain (1-200) is compared to the rest of the intrinsically disordered region (201-599) and the conserved eco1 domain (600-840). **B**. **ESCO1 N-terminus (E1N) 1-200 and neutralized mutants were fused to GFP and expressed in cells.** Diagram of ESCO1 1-200 WT and series of increasingly neutralized mutants with mutations of lysine or arginine to alanine shown by black bars on top. The PI, fraction of basic residues (FBR), and propensity for disorder calculated by PrDOS (0-0.5 = ordered, 0.5-1 = disordered) is listed for each N-terminus (without GFP) (54). **C**. **Charge-dependent chromatin association.** Confocal live cell images of HeLa cells induced to express the GFP-tagged ESCO1 N-termini shown in B. ESCO1 constructs are shown in green; DNA was stained by SiR-DNA, shown in magenta. Interphase cells are shown with a single z-slice; mitotics are shown with max intensity projections from 10 z-slices. **D**. **Quantification of interphase cells:** The experiment shown in C was repeated 3 times, and a single replicate is shown with the calculated proportion of ESCO1 in nuclei compared to the cytoplasm. **E**. **Quantification of mitotic cells:** The experiment shown in C was repeated 3 times, and a single replicate is shown with the calculated amount of ESCO1 on mitotic chromosomes compared to the cytoplasm. For D and E, (**** = p < 0.0001; *** = p < 0.001; Kruskal-Wallis test with Dunn’s correction for multiple comparisons). **F. Charge-dependent insolubility.** Flow analysis of HeLa cell lines induced to express the GFP-tagged ESCO1 peptides. Cells were fixed to show total GFP signal (green) or pre-extracted to release soluble protein and then fixed (magenta). **G**. **Quantification:** The experiment shown in F was repeated 3 times, and in each replicate the mean intensity of GFP remaining afer extraction as a proportion of the mean unextracted intensity was normalized to the WT (**** = p < 0.0001; * = p < 0.05; one way ANOVA with Tukey’s correction for multiple comparisons).

**Table 1.** Conservation of protein sequence among vertebrates. Shown are statistics for alignment among ESCO1 proteins from 53 species with human ESCO1.

| ***hESCO1 residues*** | ***Full length*** | ***1-500*** | ***1-200*** |
| --- | --- | --- | --- |
| *Pairwise similarity (%)* | 59.7 | 48.5 | 42.6 |
| *Pairwise identity (%)* | 50.1 | 37.2 | 31.6 |
| *Identity (%)* | 10.6 | 1.4 | 1.09 |
| *Isoelectric point (mean)* | 9.48 | 9.84 | 10.8 |

To test the model that the N-terminal tail of ESCO1 associates with chromatin in a charge-dependent manner, we fused the first 200 amino acids of human ESCO1 to GFP, and stably expressed this fusion in HeLa cells. As predicted, this fusion protein showed striking association with nuclei in interphase, and with condensed chromosomes in mitotic cells (*Suppl. Fig. 1A*). We then designed a series of derivatives of this construct in which the ESCO1 tail was increasingly neutralized, which we named Neut1, Neut2 and Neut 3 (Fig. 2B; *Suppl. Fig. 1B*). When similarly expressed, these derivates showed a decrease in association with chromatin, both interphase and in mitotic cells. Notably, the change in localization correlated with the amount of neutralization and could readily be detected with live cell imaging (Fig. 2C). To score this change in localization, we measured the ratio of chromatin-associated and soluble protein in interphase and mitotic cells, using the SiR DNA stain as a counterstain for chromatin (Fig. 2D and 2E). The results indicate that the fully charged wild type sequence promoted nearly complete partitioning of the fusion protein to chromatin, while even the moderately altered Neut1 mutant showed significant loss of chromatin association. The final two mutants, Neut2 and Neut3, showed almost no enrichment on chromatin.

We also assessed the impact of these neutralizing mutations in the ESCO1 tail-GFP fusions using flow cytometry. To do this, we measured remaining fluorescence after detergent extraction to remove soluble protein. Again, each increasingly neutralized mutant showed less GFP after extraction (Fig. 2F and 2G). This appeared consistent throughout the phases of the cell cycle (*Suppl. Fig. 1C*). We conclude that the localization and retention of the ESCO1 N-terminal tail depends on high positive charge.

### The N-terminal tail of ESCO1 binds DNA in a charge-dependent manner

Several lines of evidence suggest that ESCO1 may directly bind DNA. First, ESCO1 remains bound to mitotic chromosomes although Cohesin is largely evicted from mitotic chromosomes, suggesting that interaction with Cohesin is not the key factor promoting chromatin binding (13). The yeast Eco1 protein has a DNA-binding zinc-finger domain which is essential for its chromatin localization and role in Cohesin regulation, but it is dispensable in ESCO1 indicating that DNA-binding activity may be functionally conserved in a different part of the protein (13, 25). The association of ESCO1 depends on its N-terminal tail, and we have shown that it is released upon DNase treatment. As the N-terminal tail of ESCO1 is positively charged and interacts with chromatin in a charge-dependent manner we hypothesized that it is possible that the N terminal tail of ESCO1 may have a natural affinity for negatively charged DNA.

We tested whether the N terminal tail of ESCO can bind directly to DNA using and electrophoretic mobility shift assay (EMSA). We purified a recombinant N-terminal fragment of ESCO1 fused to GFP (ESCO1 1-200). Holding a ^32^P-radiolabeled single-stranded 60 base primer at a constant concentration, we titrated in the ESCO1 fusion (26, 27). Consistent with direct binding of ESCO1 to the DNA, the fusion protein resulted in greatly reduced mobility of the DNA fragment (Fig. 3A). Fitting the fraction of DNA shifted to a nonlinear curve the wildtype fusion protein gave an apparent kD of 21 nM in this assay, suggesting a relatively tight binding. To determine whether direct binding of the ESCO1 tail to DNA is charge dependent, we also tested the series of charge neutralization mutants shown in Figure 2B in the same EMSA assay. (28)As predicted by our model, the neutralized mutants showed greatly reduced ability to shift the DNA (Fig. 3C and 3D). We did note, with the exception of Neut1 that the samples bound by ESCO1 did not fully enter the gel. Retention of the protein-DNA species within the well may indicate higher-order structures or aggregates (27). Taken together, our data indicate that the human ESCO1 protein directly binds DNA through the distributed positive charge in its N-terminal tail, consistent with the data from live cell imaging shown in Figure 2.

**Figure 3.**
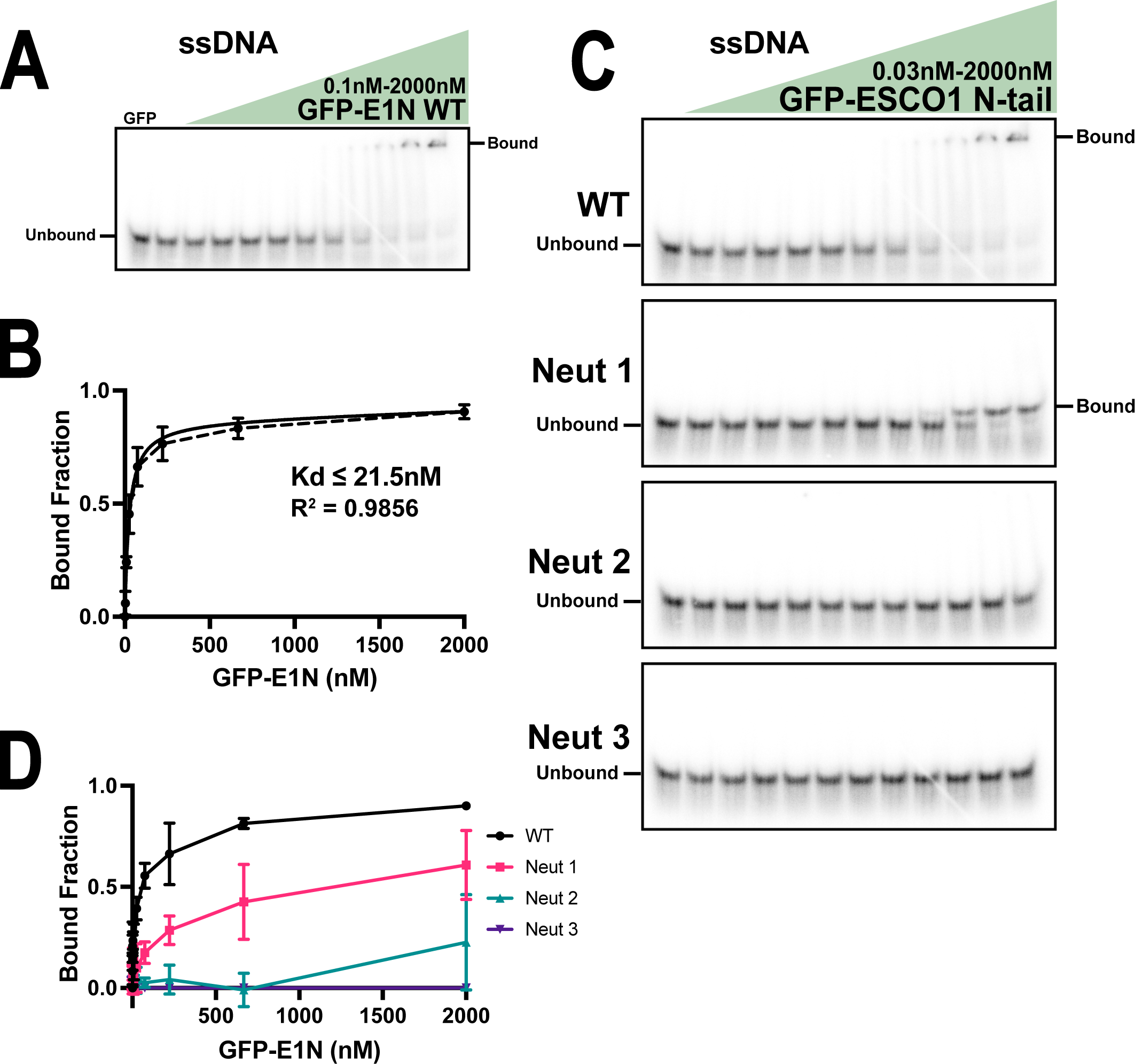
ESCO1 binds nucleic acids in vitro. A. N-terminus of ESCO1 binds DNA. An electrophoretic mobility shif assay (EMSA) using 1nM of radiolabeled 60mer ssDNA mixed with a titrating concentration (three-fold serial dilution) of GFP-E1N and ran on a native polyacrylamide gel. **B. Quantification:** The EMSA shown in A was repeated 3 times (dashed line shows the mean and standard deviation) and the data was fit to a nonlinear curve (shown as the solid black line) to calculate the apparent binding affinity (Kd). **C. Charge dependent DNA binding.** An EMSA using 1nM of radiolabeled 60mer ssDNA mixed with a titrating concentration (three-fold serial dilution) of GFP-E1N WT or neutralized mutants and ran on a native polyacrylamide gel. **D. Quantification:** The EMSAs shown in C were repeated 3 times and bound fraction was ploXed as in B.

As the primer used in the assay shown in Figure 3 was single stranded, we thought it important to also test the ability of the N terminus of ESCO1 to bind double stranded DNA. To do this, we annealed the same labeled primer to an unlabeled complementary primer and performed the EMSA again. Here, we found the same charge dependent binding of ESCO1 to dsDNA, with an apparent kD of 3.5 nM (*Suppl. Fig. 2*). The WT tail appeared to bind preferentially to longer DNA oligos over shorter (*Suppl. Fig. 3*).

### Modification of chromatin disrupts ESCO1 localization and solubility

We have shown that by neutralizing ESCO1, we reduce its ability to interact with chromatin and bind DNA. To further validate that the interaction between the ESCO1 N-terminal tail and chromatin is driven by charge, we sought to disrupt the interaction by modifying the chromatin in a manner that would alter its overall charge. Treatment of cells with trichostatin A (TSA), a deacetylase inhibitor, leads to the accumulation of acetyl groups on chromatin, most likely on histone tails. The resulting hyperacetylation of chromatin disrupts the overall chromatin state and can alter charge-dependent interactions with chromatin (23). While chromatin is usually permissive to interaction with positively charged molecules and generally excludes negatively charged molecules, hyperacetylated chromatin induced by TSA appears significantly more accessible to negatively charged proteins (23). We therefore tested whether the positively charged ESCO1 tail might show reduced interaction with TSA-treated chromatin compared to untreated controls. We imaged HeLa cells stably expressing the GFP-fused ESCO1 N-terminal tail live after treatment with 5uM TSA or an equivalent amount of DMSO vehicle as a control for 3hrs (Fig. 4A). There was a slight change in its partitioning of the fusion protein in interphase cells treated with TSA (Fig. 4B), but we found no clear effect in mitotic cells (Fig. 4C). We next tested the effect of TSA on the solubility of this ESCO1 peptide and found an increase as observed by flow analysis (Fig. 4D). Again, we saw a trending very slight difference (p = 0.0745) in the remaining fluorescence of the control and TSA-treated samples (average of 92.3% that of DMSO from three independent experiments) after extraction of soluble protein (Fig. 4E). Together, this shows that treatment with TSA and subsequent hyperacetylation of chromatin slightly disrupts the association of ESCO1 with chromatin, although much less than induced by direct neutralization of the ESCO1 protein tail. This may reflect the fact that neutralization mutations in ESCO1 cause a greater proportional change to its overall charge than TSA causes to chromatin. Alternatively, these data may indicate that positively charged proteins like ESCO1 can still bind hyperacetylated chromatin and are less affected than negatively charged proteins (23).

**Figure 4.**
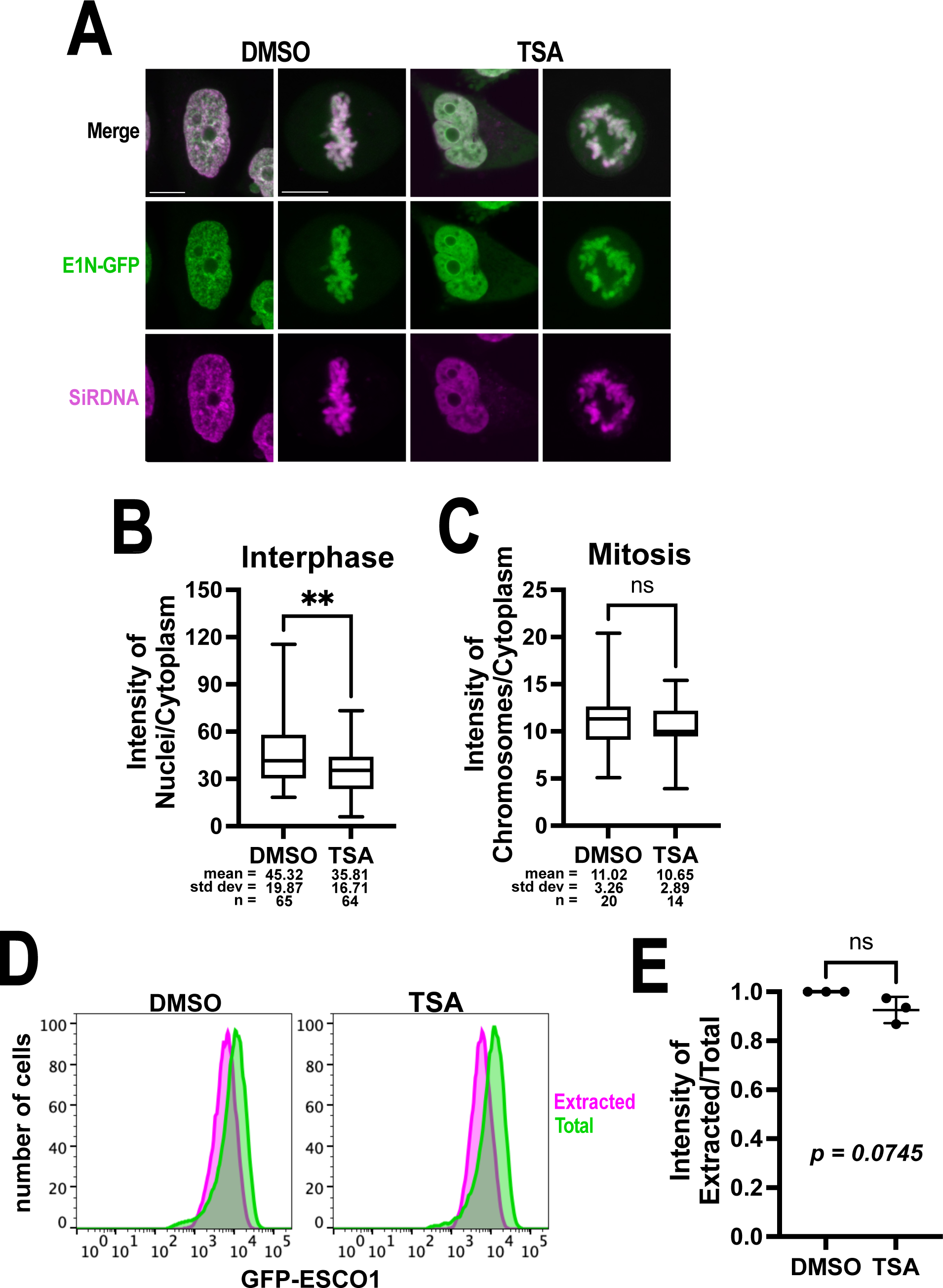
Chroma/n state has minimal effect on ESCO1 associa/on. A. Chroma/n state dependent localiza/on of ESCO1. Confocal live cell images of HeLa cells induced to express GFP-E1N WT (shown in green). Cells were treated with 5uM TSA for 3hrs to induce histone hyperacetylaJon or an equal volume DMSO as control; DNA stained with SiR-DNA (shown in magenta). **B. Quan/fica/on of interphase cells:** Experiment shown in A was repeated three Jme, and one replicate is shown with the proporJon of ESCO1 in nuclei compared to cytoplasm. **C. Quan/fica/on of mito/c cells:** Experiment shown in A was repeated three Jme, and one replicate is shown with the proporJon of ESCO1 on mitoJc chromosomes compared to cytoplasm. **D. The small change in nuclear localiza/on was consistent.** All three interphase replicates are shown, normalized to the control DMSO-treated samples. For B, C, and D, (** = p < 0.01; ns = p > 0.05; t-test). **E. Insoluble ESCO1 has minimal dependence on chroma/n state.** Flow analysis of HeLa cells induced to express GFP-E1N. Cells were fixed to show total GFP signal (green) or pre-extracted to release soluble protein and then fixed (magenta). **F. Quan/fica/on:** The experiment shown in E was repeated 3 Jmes, and in each replicate the mean intensity of GFP remaining a[er extracJon as a proporJon of the mean unextracted intensity was normalized to the control (ns = p > 0.05; t-test).

### Cohesin acetylation and sister chromatid cohesion do not depend on the ESCO1 N-terminal tail

We wondered whether the interaction of ESCO1 with chromatin through its N-terminal tail is needed for proper regulation of Cohesin. Approximately 80% of Cohesin acetylation is dependent on ESCO1, while acetylation by ESCO2 is essential for sister chromatid cohesion (10). Despite a low level of ESCO1 in the early embryos of the mouse, fish, and frog, sister chromatid cohesion occurs normally in these systems (28–30). Furthermore, sister chromatid cohesion reconstituted in Xenopus egg extracts immunodepleted of the Xenopus ESCO2 homolog, xESCO2 (also called xEco2), is rescued fully by recombinant xESCO2, but not by xESCO1 (11). In somatic human cells lacking ESCO1, sister chromatid cohesion occurs normally (10). However, upon siRNA depletion of ESCO2, cells expressing ESCO1 have improved sister chromatid cohesion compared to cells that do not express ESCO1, suggesting that ESCO1 can support minimal cohesion established by ESCO2 (10). Thus, ESCO1 acetylates Cohesin and has a supportive role in sister chromatid cohesion that is measurable upon sensitizing the system by depletion of ESCO2.

To test whether ESCO1’s interaction with chromatin through its N-terminal tail is needed for regulating Cohesin, we measured SMC3 acetylation and sister chromatid cohesion in HeLa cells lacking endogenous ESCO1, and stably expressing either flag-tagged ESCO1 WT or Δ1-200 transgenes under the control of a tetracycline-inducible promoter. We then performed siRNA-mediated knockdown of ESCO2 and measured whether expression of the ESCO1 transgene could rescue sister chromatid cohesion when ESCO2 levels were greatly reduced, as previously (10). Mitotic chromosome spreads were prepared and sister chromatid cohesion was scored as previously. As previously, we found ∼25% rescue of sister chromatid cohesion by expression of wildtype ESCO1 following siRNA depletion of ESCO2, and strikingly that the mutant ESCO1 Δ1-200 provided a similar amount of suppression of the loss of cohesion phenotype (Fig. 5C). Immunoblot analysis of the cells after 36 hours of siESCO2 and 60 total hours of Flag-ESCO1 induction showed that ESCO1 Δ1-200 was capable of acetylating SMC3 at a level comparable to WT (Fig. 5A and 5B). These experiments suggest that the direct interaction of ESCO1 with chromatin through its N terminal tail is dispensable for total SMC3 acetylation and its ability to support sister chromatid cohesion.

**Figure 5.**
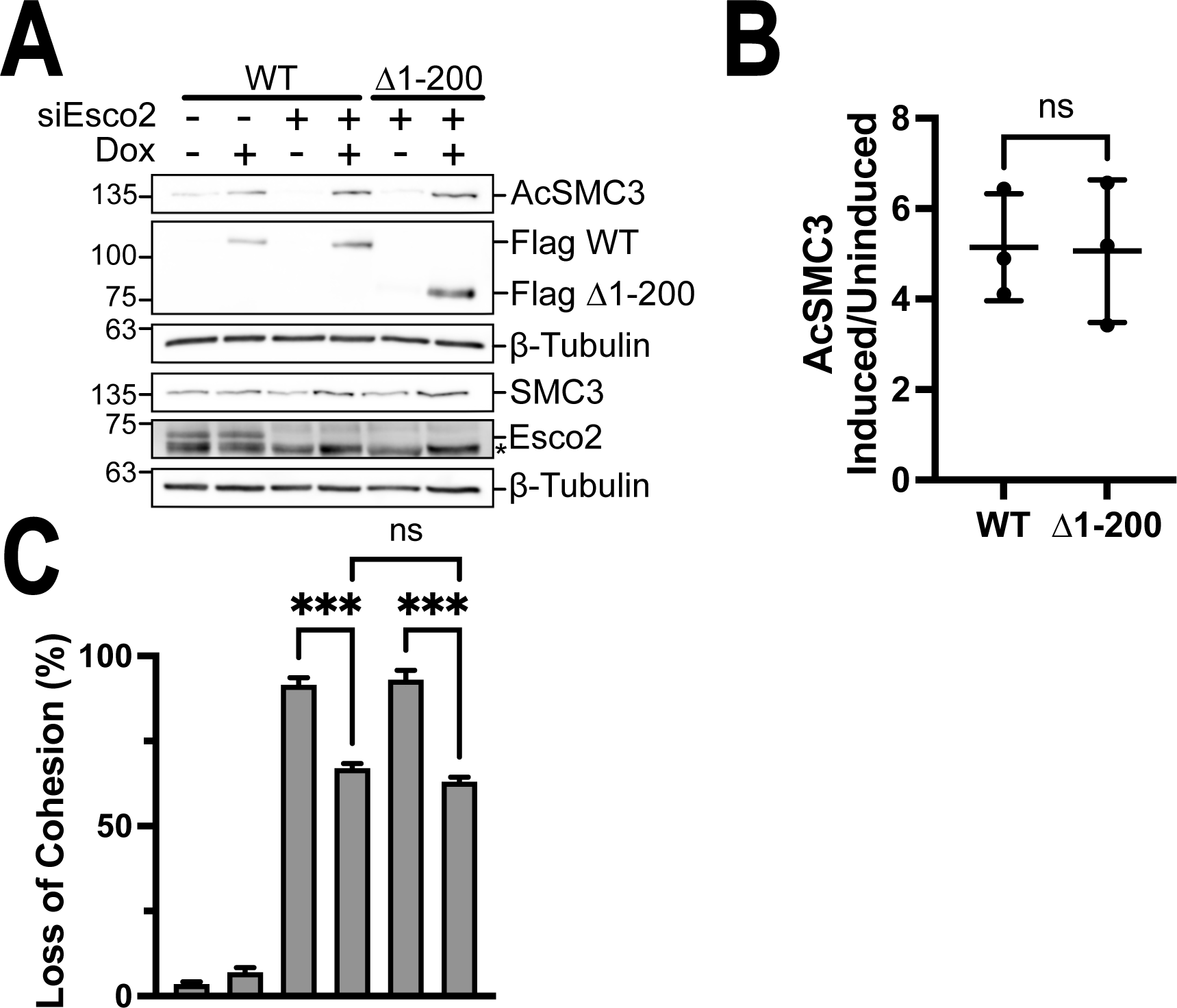
ESCO1’s N-terminal tail is dispensable for Cohesin acetylation and sister chromatid cohesion. **A. Total SMC3 acetylation is unaffected by loss of chromatin associated ESCO1**. SMC3 acetylation was analyzed by immunoblot using HeLa cells expressing flag-tagged ESCO1 WT or Δ1-200 in an siESCO2 background. **B. Quantification:** The experiment shown in A was repeated three times, signals were normalized to β-tubulin, and SMC3 acetylation was normalized to total SMC3 (ns = p > 0.05; t-test). **C. ESCO1 maintains ability to partially support sister chromatid cohesion without N-terminal chromatin association.** The experiment shown in A was repeated three times and each Dme metaphase chromosome spreads were analyzed for sister chromatid cohesion defects. Data are presented as the percentage of mitotic spreads with lost cohesion, n ≥ 100 for each sample (*** = p < 0.001; ns = p > 0.05; one way ANOVA with Tukey’s correction for multiple comparisons).

## Discussion

ESCO1 has broad functions in DNA organization, replication, repair, gene expression, and sister chromatid cohesion. How ESCO1 associates with chromosomes through its N-terminal tail, and whether this association is necessary for its ability to acetylate Cohesin and support sister chromatid cohesion was not fully understood when we undertook these studies.

We investigated how ESCO1 interacts with chromatin, both within cells and in vitro. We showed that localization and chromatin binding by ESCO1 in cells is dependent upon positive charge distributed throughout its N-terminal tail. We also demonstrated in vitro DNA binding activity by the ESCO1 N-terminal tail. This DNA binding activity, dependent on its positive charge, and seems to be the main feature ensuring ESCO1 onto chromatin. Surprisingly, we show that DNA binding by ESCO1 is dispensable for acetylating Cohesin and supporting sister chromatid cohesion (Fig. 5). This indicates that ESCO1-mediated acetylation occurs independently of its tight interaction with chromatin.

ESCO1 and ESCO2 both acetylate Cohesin and seem to support different modes of cohesion. Disruption of any of ESCO2’s interactions with replication proteins that promote its association with chromatin also disrupts it ability to acetylate Cohesin and stabilize sister chromatid cohesion (16). Here we show the opposite phenomenon for ESCO1: the primary interaction of ESCO1 with chromatin is separable from its role in acetylation of Cohesin and ability to support sister chromatid cohesion. This is particularly surprising given that multiple lines of evidence suggest that ESCO1 associates with Cohesin on chromatin. Unlike ESCO2, ESCO1 colocalizes with Cohesin in ChIP-Seq experiments, indicating that there is a prolonged or at least frequent interaction with Cohesin (9). Additionally, sequence motifs in ESCO1 that promote its colocalization with Cohesin are required for its role supporting sister chromatid cohesion but not acetylation, indicating that colocalization of ESCO1 with Cohesin, and not acetylation alone, is required for ESCO1’s role in chromosome cohesion (9, 18). 3. ESCO1 associates with a population of stably bound Cohesin-SA1 complexes throughout interphase where it is suggested to protect these Cohesins from WAPL-mediated release for hours (12).

It is possible that the small amount of ESCO1 that remains associated with chromatin in the when it is lacking its DNA-binding tail is sufficient to promote full function. A motif in the region ∼100 amino acids upstream of the DNA-binding tail described here has been shown to interact with Pds5 and is necessary for ESCO1’s roles in Cohesin acetylation and supporting sister chromatid cohesion (18). That motif and a conserved part of the catalytic domain was also predicted to interact with Pds5 (19). If ESCO1 requires association with chromatin to acetylate Cohesin, it is possible that these interactions with Pds5 or some other interaction is sufficient to promote localization of ESCO1 on chromatin with Cohesin.

Previous studies indicate that ESCO1-mediated acetylation of Cohesin depends on an interaction between ESCO1 and Pds5, results in the prevention of WAPL-mediated release of a stable population of Cohesin SA1, and underlies the presence of long-lived chromatin loops (12, 18). This would presumably be dependent upon a prolonged ESCO1 residency. However, recent studies describe a cohesin loading cycle and specifically an ESCO1-mediated acetylation cycle which regulates the binding of Pds5 to Cohesin and pausing of loop extrusion at CTCF sites (8, 31). This model of Cohesin acetylation and loading cycles would seemingly depend on an equally dynamic ESCO1. It suggests that ESCO1’s response time would affect its function in these cycles. We have shown here that significant reduction of chromatin associated ESCO1 does not reduce the total amount of acetylated Cohesin. This suggests that, in our system, chromatin associated ESCO1 is not a limiting factor in these cycles overall, but we did not investigate any genomic-site-specific effects.

ESCO1 mutants defective in supporting sister chromatid cohesion can still acetylate Cohesin, indicating acetylation is not sufficient for cohesion, as shown in the Xenopus model (9, 30). It remains to be seen whether acetylation alone is sufficient to stabilize Cohesin-mediated loops. Future studies should investigate whether this independent interaction between ESCO1 and DNA is important for Cohesin localization or loop formation. Although this is the first report of a human Cohesin acetyltransferase binding DNA, multiple Cohesin-related proteins bind nucleic acids: Pds5, SA1, SA2, and CTCF (32–37). Nucleic acid binding by SA subunits and CTCF direct Cohesin localization and chromatin organization (34, 38–41). As ESCO1 promotes Cohesin stabilization at specific sites, it seems plausible that DNA binding may help direct where and therefore contribute to organization (8, 9, 12, 18). Other proteins like ESCO1 with charged intrinsically disordered regions use them to accelerate their localization and target search (42). Since we found total Cohesin acetylation and sister chromatid cohesion was not disrupted by loss of this DNA binding tail, if there are differences in function, it would be site-specific changes in acetylation or organization that would best be observed with sequencing-based methods.

ESCO1 appears to silence expression of many genes independent of Cohesin and may organize chromatin compartments independent of Cohesin (9, 12). ESCO1’s direct interaction with DNA may play a role and should be investigated in Cohesin-dependent and potential - independent activities.

ESCO1 has a intrinsically disordered region (IDR) which contains a high concentration of charged residues. Here we show that this positively charged IDR drives ESCO1 onto chromatin through a charge-dependent interaction with DNA, consistent with recent studies which show that charge can affect partitioning of proteins to mitotic chromatin (23). The characterization of the interaction between the IDR in ESCO1 and chromatin may be relevant to similar chromatin-associated IDRs. As mitotic chromosomes are accessible to many chromatin structural proteins, these findings posit the question: how many other chromatin associated proteins associate with chromatin by charge alone (43)?

Charged and disordered protein regions comparable to the ESCO1 N-terminal tail have been shown to form phase separated droplets with DNA and condense chromatin in vitro (44, 45). In particular, linker histone has long been investigated for the role of its DNA binding C-terminal tail in driving chromatin condensation and perturbing chromatin properties (46–49). The preference of ESCO1 to bind to longer nucleic acid chains (*Suppl. Fig. 3*), is similar to a property of FUS, an RNA-binding protein that aggregates with RNAs, suggest the possibility that ESCO1 may promote crosslinking of DNA (50). Interestingly, this model is consistent with the initial identification of human ESCO1 which described protein sequence similarity to a linker histone species in tetrahymena and suggests investigations into the ability for ESCO1 to condense chromatin through this DNA binding tail (51).

## Materials and Methods

### Cell culture and stable cell lines

293T cells were obtained from the American Type Culture Collection (ATCC) and HeLa Flp-In T-Rex cells were obtained from Invitrogen. All cell lines were cultured in Dulbecco’s modified Eagle’s medium (DMEM, Corning) supplemented with 10% fetal bovine serum (FBS) (Atlanta Biologicals) and were maintained at 37 °C in a 5% CO_2_ atmosphere. Stable cell lines with doxycycline-inducible ESCO1 transgenes were generated using HeLa Flp-In T-Rex cells with endogenous ESCO1 CRISPR-knocked out as previously described (Alomer, 2017). The ESCO1 cDNAs were cloned into a pcDNA5/FRT-based C-terminally Flag- or GFP-tagged vector and co-transfected with a plasmid expressing the FLP recombinase (pOG44, Invitrogen) using Lipofectamine 3000 (Invitrogen) into Flp-In T-Rex HeLas (Invitrogen). Cells were transfected according to manufacturer’s instructions using Lipofectamine 3000 (Invitrogen) for plasmid DNA or Lipofectamine RNAiMAX (Invitrogen) for siRNA against ESCO2. Cells were selected in 200µg/mL hygromycin B (Gold Biotechnology), colonies isolated with trypsin-EDTA-soaked filter papers and expanded, and transgene induction was confirmed by immunoblot.

### Cohesion rescue assay

Cells were induced with 2µg/ml doxycycline to express the transgenes. siRNA-mediated depletion of endogenous ESCO2 was done with 20nM siRNA (Dharmacon, J-025788-11, target nucleotides: GUUGAUACCCUCAGGAAUU) in serum-free Opti-MEM (Gibco) 24 hours after induction. Media was replaced with DMEM with 10% FBS and supplemented with fresh 2µg/ml doxycycline 12 hours after siRNA transfection. Cells were collected 24 hours later. For collection, cells were washed with phosphate-buffered saline (PBS), harvested with trypsin-EDTA, washed with PBS again, swelled in 75mM KCl for 20min at room temperature (RT), and fixed in ice-cold 3:1 methanol:acetic acid. A fraction of each sample was collected in SDS for immunoblot during the second PBS wash step. Fixed samples were stored in -20°C until they were spun-down, refreshed with fixative twice, and then dropped on glass slides which dried overnight at RT. The slides were stained with Giemsa (VWR) according to the manufacturer’s instructions to visualize the DNA. Mitotic chromosome spreads were analyzed on a Zeiss Axioimager Z1 upright microscope with a 63x oil objective. Samples were scored blind and at least 100 mitotic cells for each sample were scored according to the degree of cohesion consistent in at least 10 chromosomes/cell.

### Flow cytometry

After 24 hours of induction with 2µg/ml doxycycline, T-Rex HeLas expressing GFP-tagged ESCO1 transgenes were collected for flow cytometry. Cells were washed in PBS, trypsinized, spun down, PBS washed and either pre-extracted with 0.1% Triton X-100 in PBS for 5 minutes on ice and then fixed or directly fixed in 4% PFA for 10 minutes at RT. Fixed cells were then washed twice with 20mM Tris, 150mM NaCl, 2% bovine serum albumin (BSA), 0.1% NaN_3_, 0.1% Triton X-100, again with PBS, then stained with 4µg/ml Hoechst 33342 for 45min, transferred to flow tubes on ice, and ran on a BD FACSCelesta (BD Biosciences). The average signal intensities were normalized to the average intensity of the WT in each of three independent experiments with at least 10,000 cells per sample, and then plotted together with statistical analysis from an ANOVA and Tukey’s multiple comparison follow-up (Fig. 2G) or t-test (Fig. 4E).

### Live-cell imaging

Stable HeLa cells that had been induced to express GFP-tagged ESCO1 transgenes for 24 hours were incubated with 250nM SIR-DNA (Cytoskeleton Inc.) and 2µg/ml doxycycline in Opti-MEM (Gibco) and kept in a 37 °C and 5% CO_2_ environment with a stage-top incubator (Tokai Hit). Images were collected with a 60x oil objective lens on a Nikon N2 Confocal microscope and analyzed using FIJI.

For localization measurements, a region of interest (ROI) was selected for a spot identified by SiRDNA as being within the nucleus in an interphase cell or chromosomes in a mitotic cell. An ROI of the same size was placed within the cell’s cytoplasm (identified by GFP and transmitted light) and mean intensities of GFP from both regions were collected. Local background was measured by a larger ROI and subtracted. Statistical analysis was performed by Kruskal-Wallis test with Dunn’s multiple comparisons test as follow-up.

### Cellular fractionation

Cells were fractionated as previously described (53). HeLa T-Rex cells (Fig. 1A) were harvested and resuspended in PBS at a concentration of 4 x 10^7^cells/ml (this concentration was kept throughout the experiment). After a sample was taken as whole cell lysate (WCL), cells were resuspended in buffer A (10mM HEPES pH7.9, 10mM KCl, 1.5mM MgCl2, 0.34M sucrose, 10% glycerol, 1mM DTT, 0.2mM PMSF, and 0.1% Triton X-100), incubated on ice for 5 minutes, and then spun down at 1300g for 4 minutes at 4 °C. An SDS sample was taken from the supernatant for the cytoplasm, and from the pellet for nuclei after it was resuspended in buffer A. Nuclei were spun down again at 1300g for 4 minutes at 4 °C (supernatant SDS sample again collected and pooled for cytoplasm), resuspended in 3mM EDTA, 0.2mM EGTA, 1mM DTT, and 0.2mM PMSF, and incubated on ice for 30min. Samples were spun down twice at 1700g for 4 minutes at 4 °C and each time a sample of the supernatant was taken in SDS and pooled for the soluble nuclear fraction. The pellet of the insoluble nuclear fraction was resuspended directly in SDS sample buffer. Samples were processed in one day and all samples analyzed in immunoblot together (WCL, nuclei, cytoplasm, insoluble and soluble nuclear fractions) came from the same cells. The amount of a-Flag signal remaining in the insoluble fraction was proportioned to the WCL fraction. To observe endogenous ESCO1, 293T cells (Fig. 1C) were fractionated the same way, but nuclear pellets were aliquoted, flash frozen in liquid nitrogen, and stored at -80 °C until use. Upon thawing, they were refreshed with buffer A and then digested with 125U Benzonase (Sigma-Aldrich), 50U DNase I, or 50U RNase A (Thermo Scientific) in 200ul containing 8 million cells at 37 °C for 1 minute. Nuclei were spun down again and continued processing as the HeLas. The 293T nuclei were collected and frozen down in advance, so only a single nuclear/cytoplasmic fraction in SDS sample was run as an immunoblot control. For each sample, the amount of a-ESCO1 signal in the insoluble nuclear fraction was ratioed to the total amount in both the insoluble and soluble nuclear fractions. Statistical analysis was performed by t-test (Fig. 1B) or ANOVA and Dunnett’s multiple comparison follow up (Fig. 1D).

### Immunoblot

Protein samples were resolved on 7-15% gradient polyacrylamide (SDS-PAGE) gels and transferred to nitrocellulose membranes via Trans Blot Turbo (Bio-Rad). Membranes were then blocked with 5% milk in Tris-buffered saline and 0.05% Tween 20 (TBST) for 45min at RT, incubated with primary antibodies diluted in 5% BSA in TBST and 0.1% NaN_3_ overnight at 4 °C with rocking, washed with TBST 3 times, and then secondary stained with 5% milk in TBST (for HRP-conjugated) or Intercept (TBS) blocking buffer (Li-Cor) supplemented with 0.2% Tween-20 (for IRDye-conjugated). Following 3 TBST washes and a TBS wash, blots were imaged on a Li-Cor Odyssey DLx (for IRDye-conjugated) or detected with chemiluminescence substrate (Li-Cor) and imaged on an Azure C600 CDD Imager (for HRP-conjugated). Protein signals were analyzed in FIJI by a fixed rectangular ROI around the band of interest. The local background for each lane was measured by ROI and subtracted. For acetylation of SMC3 (Fig. 5B), both SMC3 and acetylated SMC3 signals were normalized to b-tubulin, and then the amount of normalized acetylated SMC3 was proportioned to SMC3.

### Protein purification

ESCO1 N-terminal tail cDNAs were cloned into a pET-based N-terminally 6His- and GFP-tagged expression vector with a PreScission Protease cut-site between the 6His and GFP tags. These plasmids were transformed into C41(DE3) E. Coli (Sigma Aldrich) and protein expression was induced with 100uM IPTG overnight at 18 °C. Induced cells were harvested and lysed via sonication in 20mM Tris pH 8.0, 500mM KCl, 0.01% Zwittergent 3-10 detergent (Sigma), 5mM Imidazole, 5mM β-mercaptoethanol, 1mM PMSF, and 250ug/ml lysozyme (VWR). Lysates were incubated with ProteinDex Ni-NTA Agarose beads (Marvelgent) for 2 hours rotating at 4 °C to bind the 6His tag, resuspended in 20mM Tris pH 8.0, 500mM NaCl, and 5mM Imidazole, transferred into a column, and washed 5 times with the same buffer. Columns were washed twice with 50mM Tris pH 8.0 and 150mM NaCl and then the GFP and ESCO1 proteins were cleaved off the 6His and nickel beads by 630nM PreScission Protease in 50mM Tris pH 8.0, 150mM NaCl, and 5mM β-mercaptoethanol overnight, rotating at 4 °C. The flow-through was collected, protease was removed by incubation with Glutathione Sepharose Agarose beads (Thomas Scientific) for 1 hour rotating at 4 °C, beads were spun down, and cleaved protein was collected from the bead supernatant. Proteins were dialyzed to dilution buffer (50mM HEPES pH 8.0, 200mM NaCl, 1mM MgCl_2_, and 10% glycerol), concentrated, aliquoted, flash frozen in liquid nitrogen, and then stored at -80 °C.

### Electrophoretic Mobility Shift Assay (EMSA)

To perform the binding reaction, purified GFP and GFP-tagged ESCO1 N-terminal tail proteins were titrated in a threefold serial dilution (0.034nM-2000nM) over a constant concentration (1nM) of radiolabeled 60mer DNA (sequence TCGTTCCCGCAGCTTGCCAGACGCGATTTCATAGTGGAGGCCTCCAGCAATCTTGAACAC) in binding buffer (50mM HEPES pH 8.0, 200mM NaCl, 1mM MgCl_2_, 10% glycerol and 0.1% CHAPS). The dilutions and reactions were performed on ice in siliconized tubes to reduce sticking of the DNA or protein to the tube. To make double-stranded DNA, just before beginning the EMSA, the single-stranded DNA was hybridized with an equimolar complimentary 60mer (sequence GTGTTCAAGATTGCTGGAGGCCTCCACTATGAAATCGCGTCTGGCAAGCTGCGGGAACGA) in dilution buffer at 95 °C for 5 minutes and slow-cooled to room temperature by turning off the heat block. An extra lane of the radiolabeled single-stranded DNA not mixed with protein was run as a visual control for double-stranded formation, so the lowest protein concentration was removed and the titration for EMSAs with dsDNA ranged from 0.1nM-2000nM. EMSAs were resolved on native 6% polyacrylamide gels with TAE (40mM Tris pH 8.0, 20mM acetic acid, and 1mM EDTA) in the gel and running buffer. Gels were dried for 45min at a slow-cool to 80 °C, exposed to a phosphoscreen for 40 hours, and then imaged on a Personal Molecular Imager (Bio-Rad). To calculate the bound fraction in each lane, the amount of unbound (fully migrated) DNA was quantified in FIJI by a constant rectangular ROI, and local background was also measured by ROI and subtracted. The unbound signal of each lane was proportioned to the amount in the no protein control and then subtracted from 1 to get the bound fraction. The bound fraction for the series of protein concentrations from 0-2000nM was fit to a nonlinear one-site binding curve to calculate the apparent binding affinity (Kd) on PRISM.

### Protein sequence analysis and alignment – still need to make

For the ESCO1 protein sequence analysis in Figure 2A, The PI and fraction of basic residues (FBR) in ESCO1 were calculated with the aid of Lasergene (DNASTAR) and propensity for disorder was calculated by PrDOS (0-0.5 = ordered, 0.5-1 = disordered). All values and calculations made are about ESCO1 sequence and do not include consideration from any tags like GFP. The chromatin interacting domain (1-200) was analyzed compared to the rest of the intrinsically disordered region (201-599) and the conserved eco1 domain (600-840). Conservation and average PI values from ESCO1 alignments from multiple species (shown in Table 1) were generated in Geneious Prime.

### Statistical analyses

All experiments in this work were performed at least 3 times. A student or Welch’s t-test was used to compare two samples and an ANOVA or Kruskal-Wallis with multiple comparison follow-up tests were performed to compare multiple samples. For binding affinities and recovery curves, data was fit to a nonlinear least squares regression curve. PRISM v9.5 (GraphPad) was used to plot data and perform statistical analyses.

**Table 2.** Antibodies used in this study.

| **Antigen** | **Source** | **Number** | **Dilution** | **Reference** |
| --- | --- | --- | --- | --- |
| Flag | Sigma | F1804 | 1:1000 |  |
| Lamin A/C | Millipore sigma | MABT1341 | 1:5000 |  |
| β-tubulin | DSHB | E7 | 1:4000 | Chu and Klymkowsky 1989 |
| H2B | Cell Signaling | 2934 | 1:1000 |  |
| PCNA | Santa Cruz | sc-7907 | 1:1000 |  |
| ESCO1 | Lab made | OMRF168 | 1:2000 | Alomer et al. 2017 |
| SMC3 | Lab made | OMRF160 | 1:1000 | Song et al. 2010 |
| AcSMC3 | Gift from K. Shirahige |  | 1:1500 | Nishiyama et al. 2010 |
| ESCO2 | Bethyl labs | A301-689A | 1:3000 |  |
| GFP | Proteintech | 66002-1-ig | 1:5000 |  |
| GAPDH | GeneTex | GTX627408 | 1:1000 |  |
| Donkey a-mouse (HRP conjugated) | Jackson ImmunoResearch | 715035151 | 1:5000 |  |
| Goat a-rabbit (HRP conjugated) | Thermo Fisher Scientific | 31460 | 1:10000 |  |
| Goat a-mouse (IRDye 800CW) | Licor | 926-32210 | 1:20000 |  |
| Goat a-rabbit (IRDye 800CW) | Licor | 926-32211 | 1:20000 |  |
| Goat a-mouse (IRDye 680RD) | Licor | 926-68070 | 1:20000 |  |

**Table 3:** Plasmids used in this study.

| **IDENTIFIER** | **Contents** | **Source** |
| --- | --- | --- |
| **PEL06** | pcDNA5/FRT 3Flag, ESCO1 WT | This work |
| **PJRS45** | pcDNA5/FRT 3Flag, ESCO1 Δ1-200 | This work |
| **PJRS54** | pcDNA5/FRT GFP-ESCO1(1-200) neut 1 | This work |
| **PJRS55** | pcDNA5/FRT GFP-ESCO1(1-200) neut 2 | This work |
| **PJRS56** | pcDNA5/FRT GFP-ESCO1(1-200) neut 3 | This work |
| **PJRS57** | pcDNA5/FRT GFP-ESCO1(1-200) WT | This work |
| **PJC297** | pET28-6His-PreSc-GFP | This work |
| **PJC298** | pET28-6His-PreSc-GFP-ESCO1(1-203) WT | This work |
| **PJRS48** | pET28-6His-PreSc-GFP-ESCO1(1-200) neut 1 | This work |
| **PJRS49** | pET28-6His-PreSc-GFP-ESCO1(1-200) neut 2 | This work |
| **PJRS50** | pET28-6His-PreSc-GFP-ESCO1(1-200) neut 3 | This work |
| **POG44** | FLP recombinase (Invitrogen) | Invitrogen |

## Acknowledgments

We thank Katsuhiko Shirahige for the anti-acetyl SMC3 used in this study. We are grateful to the members of the Rankin lab for helpful discussions during this work. We also thank all members of the Program in Cell Cycle & Cancer Biology and Department of Cell Biology for their input and discussion during this work.

---

**Supplementary Figure 1.**
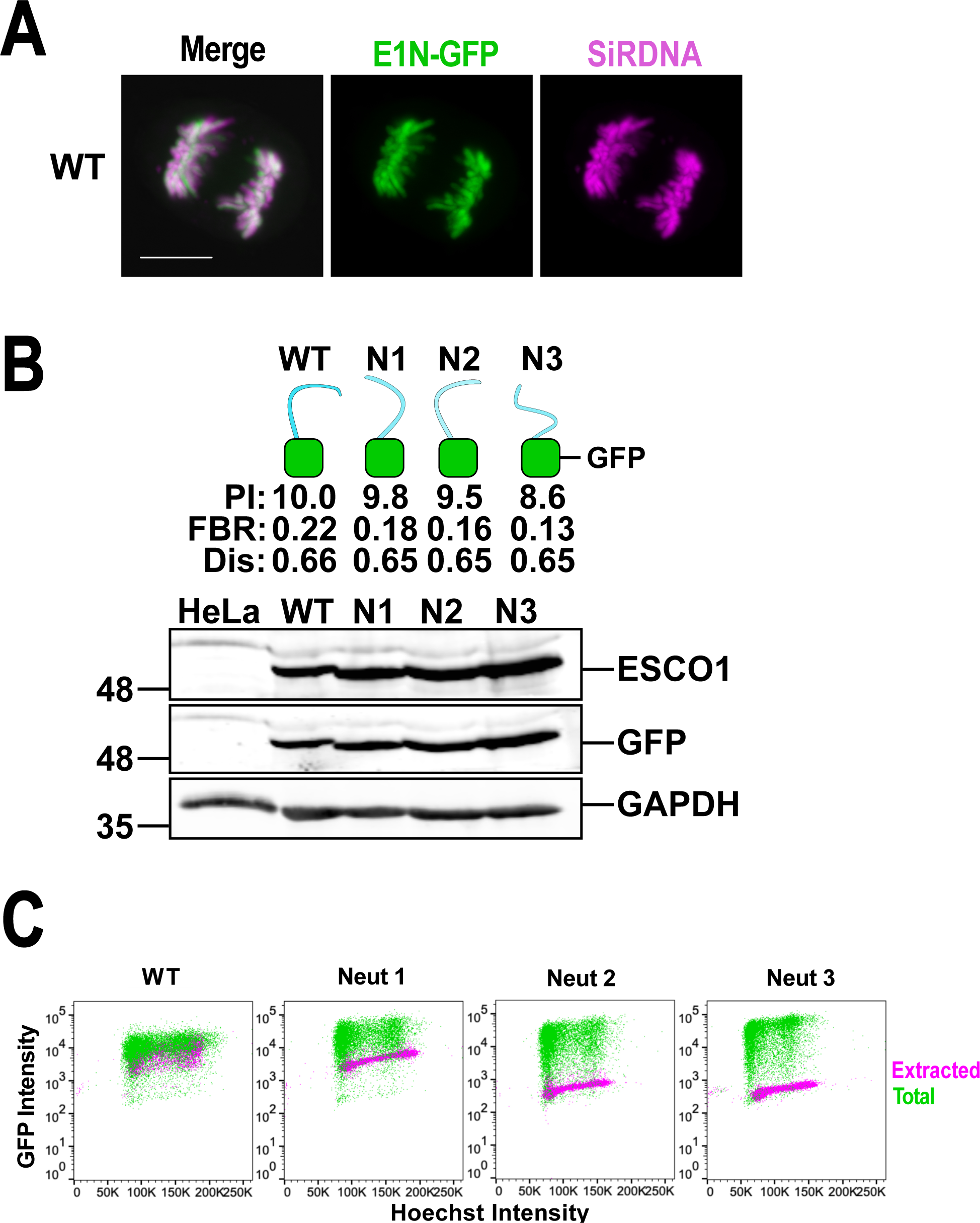
Charge drives ESCO1 1-200 fused to GFP to chromatin. A. ESCO1 1-200 (E1N) binds to condensed mitotic chromosomes. Confocal live cell image of a HeLa cell induced to express the GFP-tagged ESCO1 N-terminus. ESCO1 construct is in green; DNA was stained by SiR-DNA, shown in magenta. This anaphase cell is shown with max intensity projections from 10 z-slices. **B. ESCO1 1-200 and a series of neutralized mutants were fused to GFP and expressed in cells.** A diagram of GFP-tagged E1N WT and mutants is shown and an immunoblot validating the expression of the transgenes is shown. **C. Charge neutralized mutants are more soluble throughout the cell cycle.** The experiment in Figure 2F is shown again with GFP signal ploXed on the y-axis over the DNA signal (Hoechst 33342) on the x-axis.

**Supplementary Figure 2.**
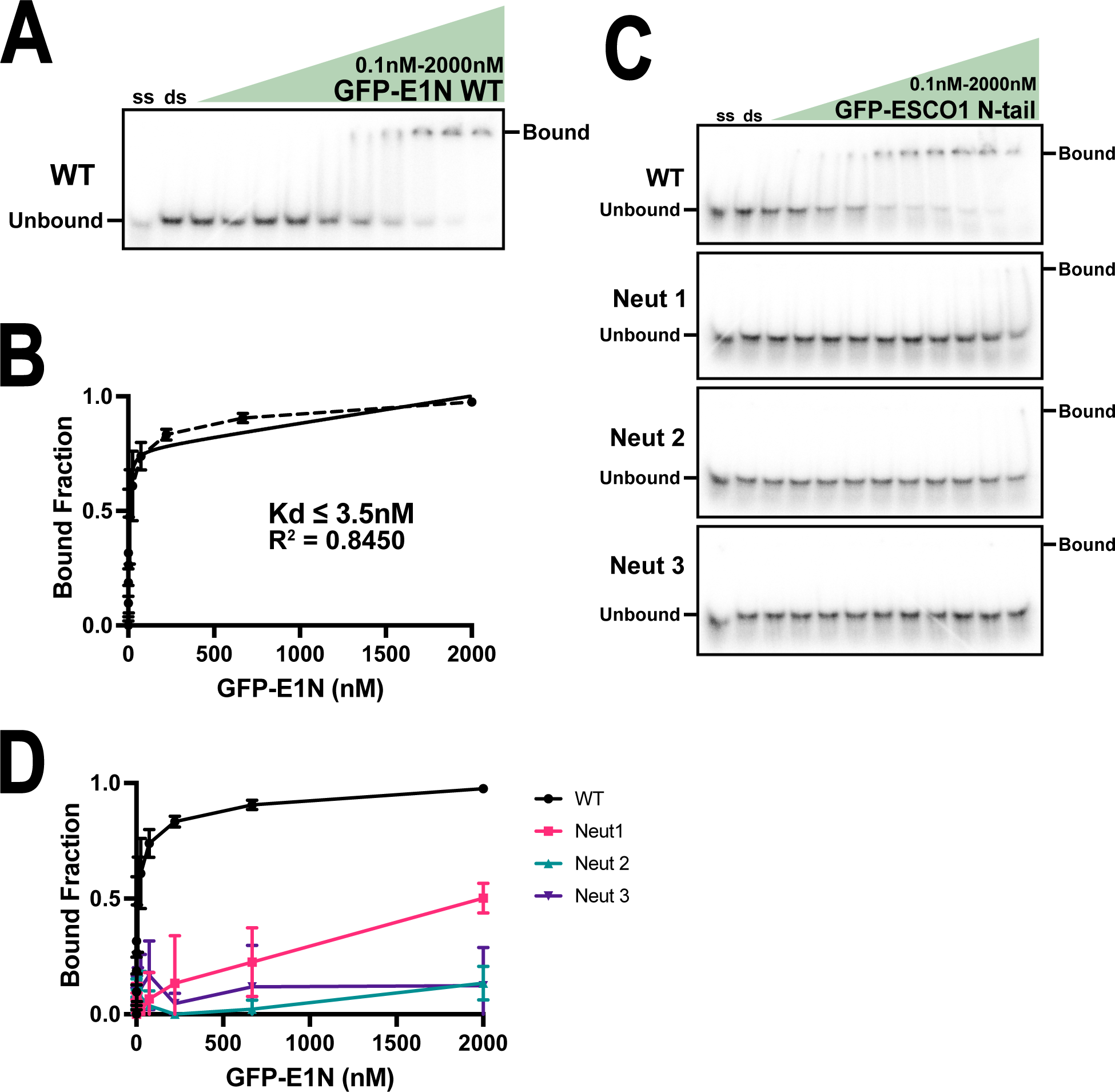
ESCO1 binds dsDNA in vitro. A. N-terminus of ESCO1 binds dsDNA. An EMSA using 1nM of radiolabeled 60mer dsDNA mixed with a titrating concentration (three-fold serial dilution) of GFP-E1N and ran on a native polyacrylamide gel. **B. Quantification:** The EMSA shown in A was repeated 3 times (dashed line shows the mean and standard deviation) and the data was fit to a nonlinear curve (shown as the solid black line) to calculate the apparent binding affinity (Kd). **C. Charge dependent dsDNA binding.** An EMSA using 1nM of radiolabeled 60mer dsDNA mixed with a titrating concentration (three-fold serial dilution) of GFP-E1N WT or neutralized mutants and ran on a native polyacrylamide gel. **D. Quantification:** The EMSA shown in C was repeated 3 times and bound fraction was ploXed as in B.

**Supplementary Figure 3.**
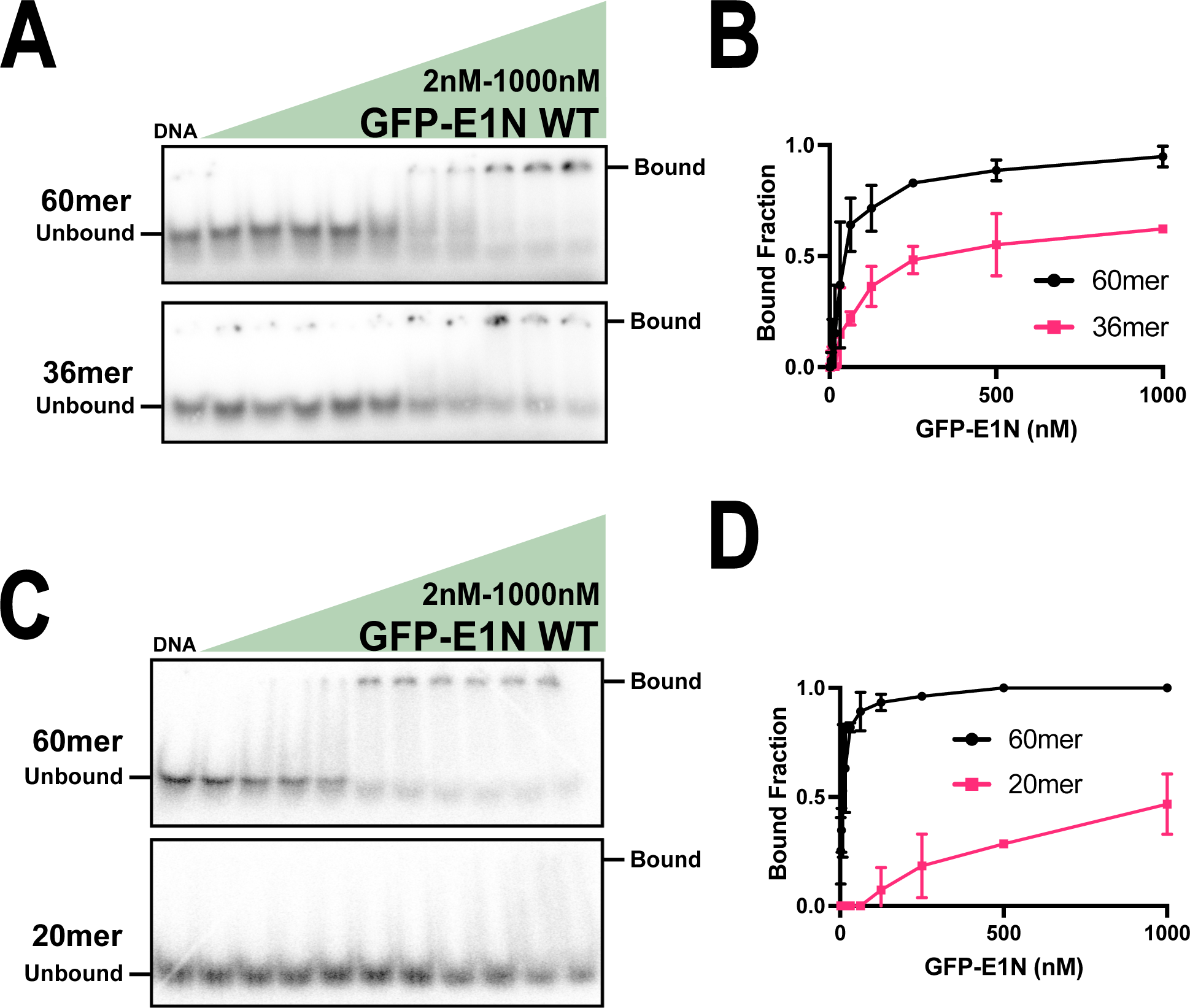
Oligo size-dependent affinity. A. ESCO1 has higher affinity for 60mer than 36mer. An EMSA using 1nM of radiolabeled 60mer or 36mer ssDNA mixed with a titrating concentration (two-fold serial dilution) of GFP-E1N and ran on a native polyacrylamide gel. **B. Quantification:** The EMSAs shown in A were repeated 2 times and the bound fraction was ploXed as a function of protein concentration. **C. ESCO1 has higher affinity for 60mer than 20mer.** An EMSA using 100pM of radiolabeled 60mer or 20mer ssDNA mixed with a titrating concentration (three-fold serial dilution) of GFP-E1N WT and ran on a native polyacrylamide gel. **D. Quantification:** The EMSAs shown in C were repeated 2 times and bound fraction was ploXed as in B.

